# A nematode model to evaluate microdeletion phenotype expression

**DOI:** 10.1101/2022.11.09.515676

**Authors:** Katianna R. Antkowiak, Peren Coskun, Sharon T. Noronha, Davide Tavella, Francesca Massi, Sean P. Ryder

## Abstract

Microdeletion syndromes are genetic diseases caused by chromosomal deletions too small to be detected by karyotyping. They are typified by complex pleiotropic developmental phenotypes that depend both on the extent of the deletion and variations in genetic background. Microdeletion alleles disrupt several genes simultaneously, often as the result of a single mutagenic event, causing a wide array of consequences across multiple systems involving multiple pathways. How simultaneous haploinsufficiency of numerous adjacent genes leads to complex and variable pleiotropic phenotypes is not well understood. CRISPR/Cas9 genome editing has been shown to induce microdeletion-like alleles at a meaningful rate. Here, we describe a microdeletion allele in *Caenorhabditis elegans* recovered during a CRISPR/Cas9 genome editing experiment. We mapped the allele to chromosome V, balanced it with a reciprocal translocation crossover suppressor, and precisely defined the breakpoint junction. The allele simultaneously removes 32 protein-coding genes, yet animals homozygous for this mutation are viable as adults. Homozygous animals display a complex phenotype including maternal effect lethality, producing polynucleated embryos that grow into uterine tumors, vulva morphogenesis defects, body wall distensions, uncoordinated movement, and a shortened life span typified by death by bursting. Our work provides an opportunity to explore the complexity and penetrance of microdeletion phenotypes in a simple genetic model system.

## INTRODUCTION

The microdeletion syndromes are complex pleiotropic developmental diseases caused by multi-locus deletions less than five megabases (Mb) in length. Microdeletion syndromes display variable penetrance depending upon the extent of the deletion and the genetic background. The most common is 22q11.2DS, also known as DiGeorge or velocardiofacial syndrome [1-4]. This disease affects an estimated one in 4000 people and manifests a wide variety of disease phenotypes, including heart abnormalities, craniofacial defects, cleft palate, short stature, psychiatric and intellectual disorders, and immunodeficiency (reviewed in [1, 2]). About 10% of cases are familial with an autosomal dominant inheritance pattern [5], and the rest result from spontaneous deletions formed during gametogenesis [6]. The extent and impact of the syndrome can vary widely among diagnosed patients, and some individuals are nearly asymptomatic. Ninety percent of cases are caused by a three megabase deletion on chromosome 22 that disrupts approximately 46 protein-coding genes. Current evidence suggests that the disease phenotypes are caused by complex genetic interactions between multiple disrupted genes and are strongly influenced by genetic background [2, 3, 6]. A pressing question, not just for this disease but more generally for all microdeletion syndromes, is whether these interdependencies can be exploited to develop therapies to mitigate disease severity. More broadly, understanding how complex multigenic phenotypes manifest will help better clarify nonlinear relationships between genotype and phenotype.

Microdeletion alleles can also arise during targeted genome editing experiments [7, 8]. Cas9 is an RNA-guided double stranded DNA endonuclease encoded by the *Streptococcus pyogenes* CRISPR locus [9, 10]. It defends the bacterial genome from phage, transposons, and other genome invaders by targeting and destroying DNA sequences derived from prior infections of the host or its ancestors. In the lab, Cas9 is widely used as a tool to generate targeted mutations through site specific DNA cleavage and repair [11-16]. Cas9 can be programmed with a hybrid guide RNA to target almost any sequence, a property that makes it fundamentally useful to genome editing [17]. The ease by which Cas9 can be programmed, coupled to its relatively high specificity, has led to the widespread adoption of this technology [11-16, 18-28]. There is also extraodinary interest in developing this technology to treat human disease (reviewed in [29]). However, experiments in mouse and human cells and embryos reveal that Cas9 cleavage can cause microdeletion lesions, leading to new concerns about applying this technique to patients [7]. The frequency at which such events happen is not well understood. On target “collateral damage”, as it has been termed, can be difficult to detect using PCR or short-read based DNA sequencing technology [8].

The small bacterivorous nematode *Caenorhabditis elegans* is widely used to study gene function in aging, development, and numerous other aspects of animal physiology. It can be grown to high yield at room temperature in solid or liquid culture in a short time. There is a robust genetic tool set to enable simple mapping of mutations recovered from genetic screens. It can reproduce as a hermaphrodite simplifying the maintenance of stocks, yet it can be crossed to facilitate creation of multiple mutant lines. The genomes of *C. elegans* and closely related species have been sequenced, and the genome size is relatively small (∼100 Mb) making whole genome re-sequencing cost-efficient. The animal is transparent, even as an embryo, enabling live imaging of organ and tissue development in real time. A large collection of mutant alleles—including large deficiencies—is widely available to researchers through the *Caenorhabditis* Genetics Center. In addition, thanks to CRISPR/Cas9, it is now routine to make targeted gene disruptions and replacements in this model organism [11, 13, 18-28].

While screening F2 progeny during the process of making a CRISPR knock-in mutation in *C. elegans*, we identified animals with an unusual uterine tumor phenotype that are prone to bursting. We mapped the causative mutation (*sprDf1*) to a ∼0.25 Mb microdeletion allele on the left arm of chromosome V that removes 32 protein-coding genes. Remarkably, homozygous animals survive to adulthood, but are small, sterile, form large uterine tumors derived from polynucleated misshapen embryos. The animals are uncoordinated and most of them die by bursting within eight days post-hatching. While many deficiency alleles have been genetically mapped, most are embryonic lethal as homozygotes and have primarily been used for mapping and assessing genetic nulls [30, 31]. Of the 212 multilocus deletion alleles described in Wormbase whose precise genomic coordinates are known, only ten are larger than 10KB, and none are over 50KB [32]. This 255KB *sprDf1* microdeletion is five times larger than the next biggest physically-mapped multilocus deletion allele in the worm. As such, it provides a unique opportunity to assess microdeletion-like genetic interactions in a simple model organism. Here, we present the initial characterization of this microdeletion allele as a potential model for the complex genetic interactions induced by similar alleles in humans.

## RESULTS

### Recovery of a homozygous viable microdeletion on chromosome V

The original goal of our work was to make a knock-in mutation of the *mex-5* locus using CRISPR-HR. We sought to recode endogenous *mex-5* to produce a variant that stabilizes the protein fold [33]. Towards that end, we designed a guide RNA that targets the *mex-5* locus on chromosome IV and a homologous recombination repair template encoding the stabilized variant. We microinjected purified recombinant Cas9 enzyme in complex with the *mex-5* guide, a guide that targets *dpy-10* as a co-CRISPR marker, and the repair template. While analyzing the progeny of injected animals, we identified sterile individuals expressing an unusual uterine tumor phenotype characterized by swelling of the body wall (**Fig. 1A**). The *mex-5* gene is essential for body axis formation during early embryonic development and null mutations are maternal effect lethal producing dead eggs [34]. Because our intended knock in mutation stabilizes the fold of MEX-5 [33], we wondered if this unexpected uterine tumor phenotype might be a neomorphic event caused by the mutation we designed. Consistent with this hypothesis, we could detect the presence of the knock in allele using a PCR and restriction digestion strategy that selectively cleaves the recoded allele (**SFig. 1**).

**Figure 1.**
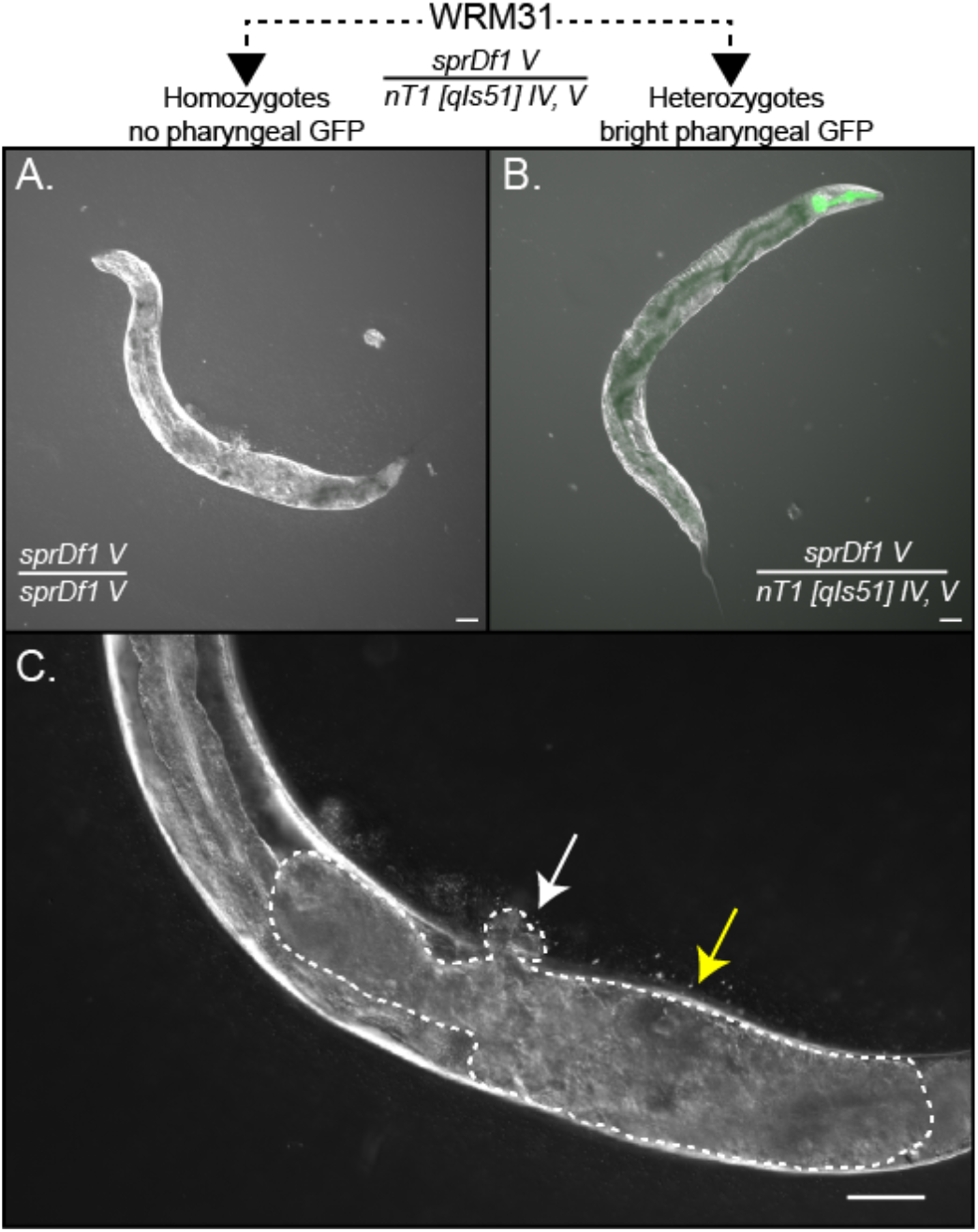
The *sprDf1* microdeletion induces a complex pleiotropic phenotype. The WRM31 strain is a heterozgous animal that contains a microdeletion allele (*sprDf1*) on chromosome V and an nT1[qIs52] balancer chromosome that supresses crossover repair of the deficiency, harbors a lethal mutation that prevents recovery of viable nT1[qIs52] homozygotes, and encodes an integrated pharyngeal GFP marker. Only two viable genotypes are produced by self-fertilization of this strain: GFP– homozygotes [*sprDf1*/*sprDf1* V] and GFP+ heterozygotes [*sprDf1* V / nT1[qIs52] IV, V]. **A-B**. Adult GFP– *sprDf1* homozygous worms are shorter and thicker than their GFP+ heterozygous siblings and form a large uterine tumor **(C)** that causes a protruding vulva (white arrow) and frequently leads to body wall distentions (yellow arrow). The phenotype is easy to follow using standard light microscopy. The scale bar in panels A and B represents a distance of 50 microns, and in panel C it represents 20 microns.

To simplify propagation, we crossed fertile siblings of the phenotypic mutants with VC362, a strain that harbors the nT1 [qIs51] IV;V reciprocal translocation chromosome that acts as an effective balancer for the *mex-5* locus. Homozygous nT1 [qIs51] IV;V animals die as larvae due to a presence of an uncharacterized lethal mutation on the balancer chromosome [35]. Heterozygous animals grow to adulthood, are fertile, and are easily identified under a fluorescence microscope through a bright pharyngeal GFP transgene integrated on the balancer chromosome [35] (**Fig 1B**). Animals homozygous for the mutation lack the balancer chromosome and thus pharyngeal GFP, simplifying identification of homozygous mutant self-progeny.

However, in the process of outcrossing this strain through wild-type N2 worms, we discovered that the unusual uterine tumor phenotype persists in animals that no longer harbor the recoded *mex-5* allele, suggesting the phenotype derives from a co-occurring mutation also balanced by nT1[qIs51]. Being intrigued by the phenotype, we decided to map the causative locus by crossing the mutant, made in the N2 background, with the divergent Hawaiian strain (CB4856) that contains numerous SNPs that facilitate recombination mapping efforts [36]. F2 progeny from the cross were singled, allowed to grow to adulthood and assessed for the uterine tumor phenotype shown in **figure 1**. We then determined the frequency of recombination in phenotypic animals using single worm PCR and a suite of DraI sensitive SNPs covered by the nT1[qIs51] balancer [36]. This analysis revealed the mutation is located to the left of snp_Y61A9LA / WBVar00208343 (recombination frequency 16.7%, position 4,550,942 on Chromosome V, **Fig. 2A**).

**Figure 2.**
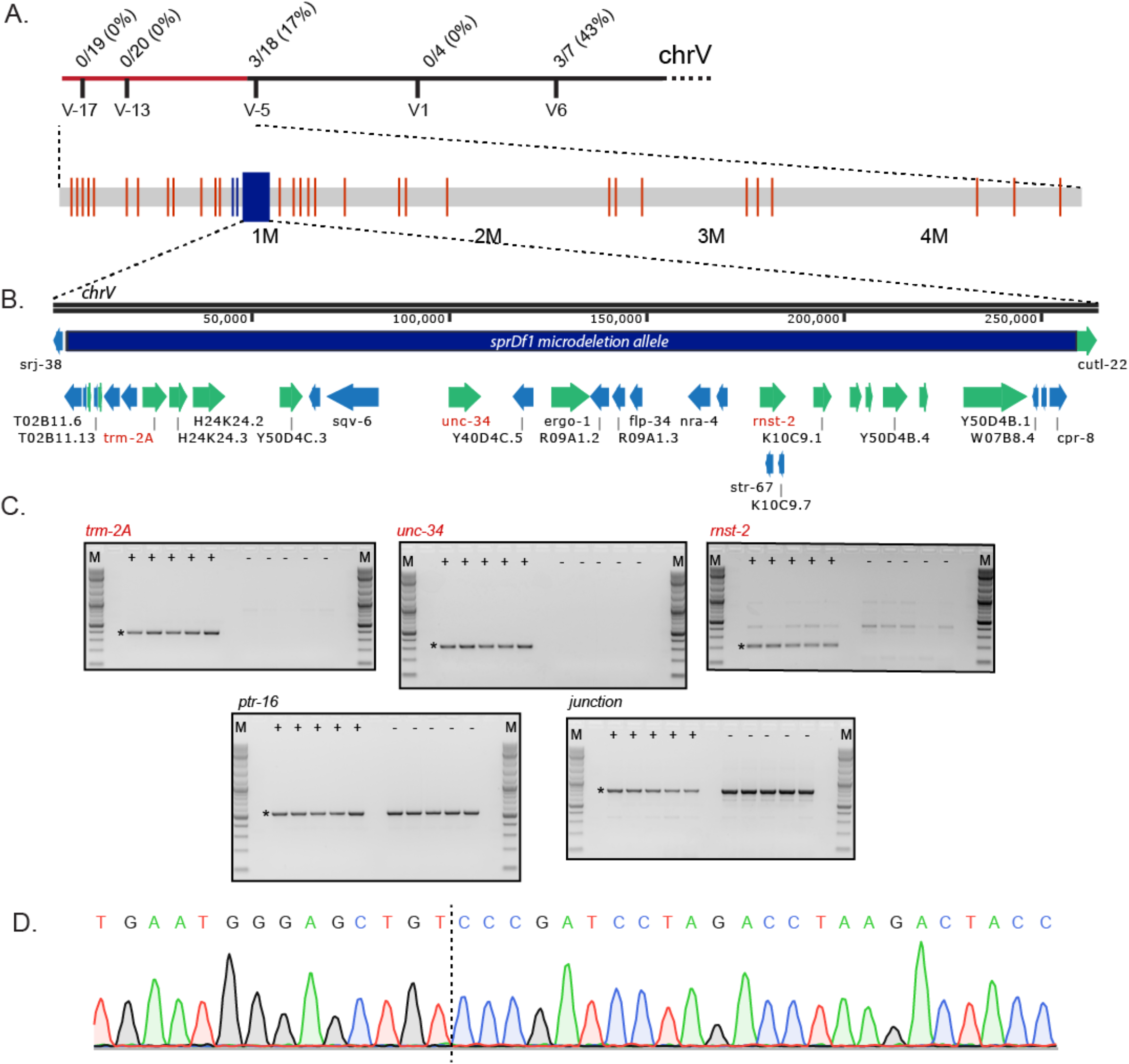
Characterization of the *sprDf1* microdeletion allele. **A**. Recombination analysis following a cross of WRM31 with the Hawaiin strain (CB4856). The number of recombination events at each SNP is represented by the numerator, while the denominator represents the number of phenotypic cross progeny assessed at each SNP. The mutation lies to the left of SNP snp_Y69A9L (V-5) on chromosome V. This region is shown below with red lines indicating the position of WRM31-specific SNPs and Indels detected by whole genome sequencing, and blue rectangles indicating WRM31 specific structural variants. **B**. The *sprDf1* microdeletion allele. The identities and positions of the protein coding genes eliminated in the microdeletion are shown below the map. **C**. Single worm PCR of multiple GFP+ and GFP– siblings (labeled) to evaluate the presence or absence of three genes predicted to be eliminated by the microdeletion (*trm-2A, unc-34*, and *rnst-2*), one gene that lies just outside of the microdeletion allele (*ptr-16*), and with primers that flank the breakpoint predicted in whole genome sequencing of WRM31. In all cases, the expected amplicon product is marked with an asterisk. **D**. Sequencing trace revealing the precise junction of the microdeletion allele (dashed line).

To further identify candidate mutations, we next performed whole genome sequencing of DNA prepped from mixed stage animals of the parental N2 strain, VC362 (the source of the nT1[qIs51] balancer chromosome), and WRM31, which harbors the balanced candidate mutation (**Fig. 1**). This analysis identified 39 SNPs or short indels unique to WRM31 that map to the left arm of chromosome V (**Fig. 2B, STable 1**). Thirty-seven have an allele frequency of ∼0.5, as expected for a heterozygous allele. Of these, 15 are intergenic, 10 are within introns, 9 are upstream or downstream of a known or predicted gene, 2 are within annotated UTRs, and 1 is in the exon of a non-coding RNA gene. None are found within a protein coding exon.

Analysis of potential structural variations identified three additional candidates unique to WRM31 on the left arm of chromosome V that perturb one or more open reading frame. The candidates are a 633 base pair deletion within K09C6.9, a 1.02 kilobase pair deletion within *srj-38*, and a 255 kilobase pair microdeletion predicted to disrupt 32 protein coding genes. To validate the presence or absence of the monogenic K09C6.9 and *srj-38* deletions in WRM31, we designed primer sets to detect the predicted deletions and performed single worm PCR comparing homozygous GFP– phenotypic animals to heterozygous GFP+ siblings. The results confirmed the absence of the deleted region in *srj-38* but not K09C6.9. This suggests that the latter is not responsible for the phenotype but leaves the deletion allele in *srj-38* as a potential candidate.

SRJ-38 is a seven transmembrane domain G-protein coupled receptor expressed in neurons and the pharynx. There are 24 mapped alleles of this gene, and 46 naturally occurring variants, none of which have been associated with a phenotype. In addition, no phenotypes have been reported upon knockdown in previous genome-wide RNAi screens. Consistent with previous findings, we observe no phenotype upon RNAi knock down of *srj-38* performed by soaking our N2 worm stock with a double stranded RNA sequence targeting this gene. This suggests the unusual uterine tumor phenotype is not likely to be caused by loss of function of the *srj-38* locus.

To test whether the large 255KB microdeletion allele is present in mutant animals as opposed to a mapping artifact, we designed primer sets to detect three genes that lie within the microdeletion (*rnst-2, unc-34*, and *trm-2A*), and another that lies just outside of the microdeletion breakpoint (*ptr-16*). PCR of single worm lysates made from heterozygous GFP+ worms robustly detect all four amplicons, while PCR of lysates made from phenotypic GFP– siblings only detect *ptr-16* (**Fig. 2C**). To further confirm the deletion allele, we designed primers flanking the predicted break point junction. These primers only amplify the deletion allele yielding a predicted product of 768 base pairs. Consistent with the presence of a large microdeletion allele, we observed an ∼800 base pair band in both GFP+ heterozygotes and GFP– homozygotes (**Fig. 2C**). We cloned and sequenced this PCR product, which identified the precise microdeletion junction. The left side break point lies within exon 6 of the T02B11.6 locus, and the right side break point lies within an intergenic region between *cpr-8* and *cutl-22* (**Fig. 2D**). In total, 255,603 base pairs are lost in this deletion, eliminating 32 characterized or predicted protein coding genes (**STable 2**).

### The microdeletion induces a uterine tumor phenotype

WRM31 GFP– animals display a wide array of phenotypes. Most notably, GFP– animals form large, disorganized tumors within the uterus after the last larval molt (**Fig. 1**). These apparent tumors grow to a large size causing distension of the body wall. To quantitatively assess these phenotypes, we cloned single L3/L4 larvae to individual plates under fluorescence optics to segregate GFP– homozygotes from GFP+ heterozygous siblings. We measured the total brood (number of embryos laid on the plate), the number of viable progeny, and the hatch rate for each animal in both genotypes (**Fig. 3A**). Of the 120 GFP– animals scored across three biological replicates, 118 were sterile, lacking the ability to deposit eggs on the plate. No viable hatchlings were retained in the uterus, though in all cases the uterus was filled with apparent mishappen embryos that grew to large size. The remaining two GFP– animals were fertile and displayed no signs of the uterine tumor phenotype. Single worm PCR revealed that these animals lost the balancer chromosome and are not mutants. Excluding these two animals, all remaining isolates displayed the uterine tumor phenotype with body wall distension shown in **figure 1**.

**Figure 3.**
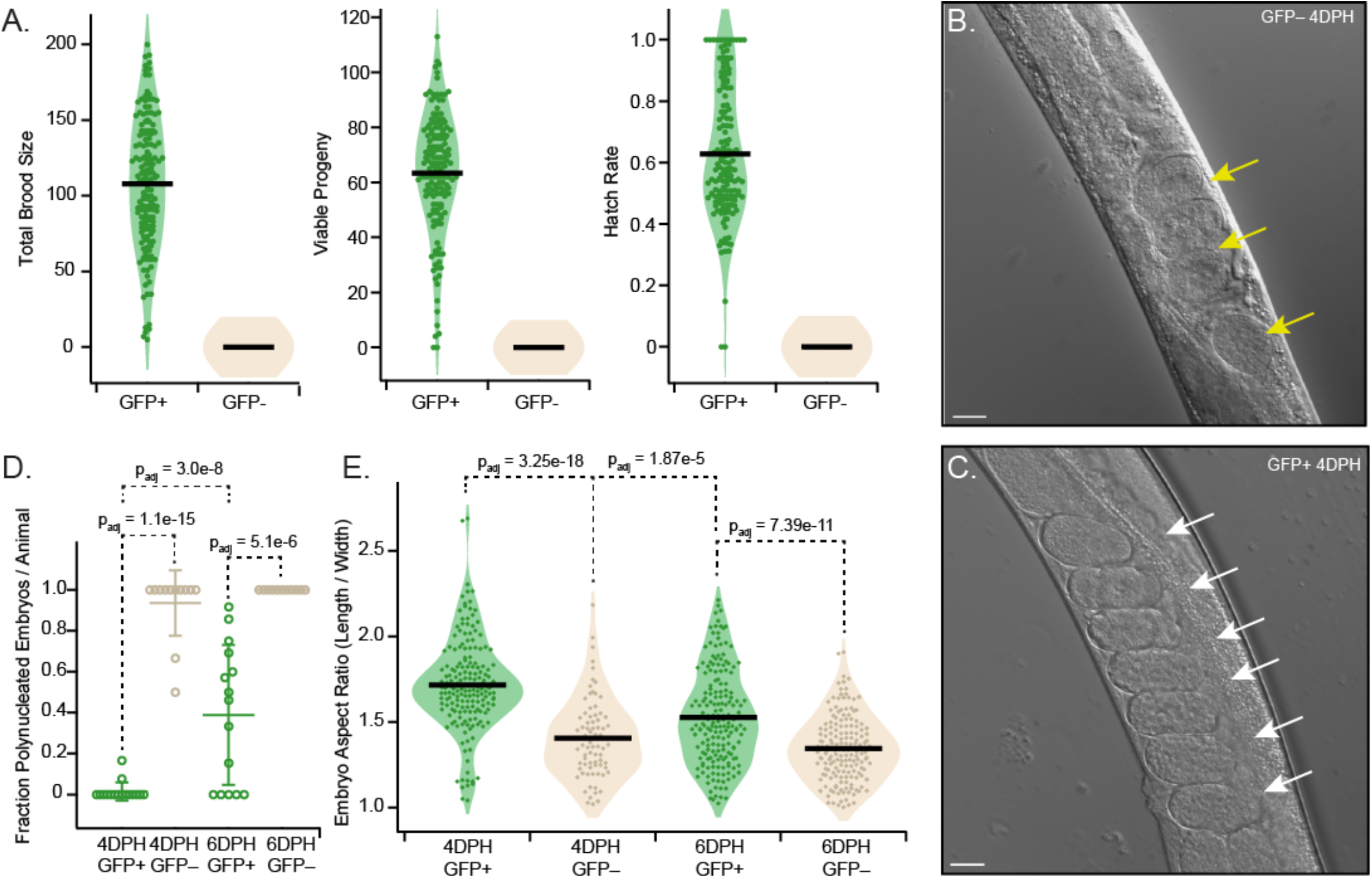
The *sprDf1* allele induces a maternal effect lethal phenotype with polynucleated misshapen embryos. **A**. The total brood size was determined by counting the number of embryos and animals deposited on a plate by a single hermaphrodite adult throughout its fertile life span. The total viable brood and hatch rate were determined by counting the number of embryos that hatched. In all panels, the green dots represent GFP+ heterozygous animals, while beige dots represent GFP– homozygous animals. The black bar denotes the mean. The hatch rate was calculated by dividing the number of viable animals by the total brood. **B-C**. Representative micrographs of young adult GFP– homozygotes (B) and GFP+ heterozygotes (C) collected four days post hatching. The yellow arrows indicate polynucleated embryos, while the white arrows represent embryos that appear normal. The scale bars represent a distance of 20 microns. **D**. The fraction of polynucleated embryos retained in utero per animal was determined at day four and six post hatching. The mean is denoted by a horizontal bar, and the standard deviation is represented by the error bars. The p-values were calculated by one-way ANOVA with post-hoc Bonferroni correction for multiple hypothesis testing. **E**. The length and width of each imaged embryo was determined from the images collected in D, and the aspect ratio (length / width) was also calculated for each embryo. The green dots represent the aspect ratio for embryos produced by GFP+ hermaphrodites, and the beige dots represent embryos produced by GFP– siblings. The black bars represent the mean, and statistical significance was assessed as in panel D.

In contrast, 153 out of 155 GFP+ animals scored were fertile and produced viable progeny, albeit at a lower rate than N2 worms. The worms produced an average of 108 ± 43 embryos and 63 ± 22 viable progeny per isolate, yielding an average hatch rate of 63 ± 21 %. Intriguingly, the brood size and hatch rate for heterozygous (GFP+) animals was quite variable even among siblings in the same biological replicate. Two worms produced a limited number of embryos that failed hatch. Several more produced less than twenty live progeny. The full distribution in both viable offspring, total brood, and hatch rate can be seen in **figure 3A**.

### Homozygous microdeletion mutants form mishappen polynucleated embryos

To better characterize the origin of the uterine tumor, we synchronized L1 animals and imaged the uterus as a function of the number of days post hatching by differential interference contrast microscopy. We observed the presence of embryos in the uterus shortly after the onset of adulthood. We imaged retained embryos at day four and day six post-hatching in both homozygotes and heterozygous siblings (**Fig. 3B**). At day four post hatching, 97% of the embryos in GFP– animals appeared polynucleated (74/76 embryos imaged across 13 animals, **Fig. 3B–D**). All of the animals imaged contained polynucleated embryos. The two embryos that were not polynucleated appear to have been recently fertilized and lacked visible nuclei. The embryos produced by GFP– animals were unusually round compared to normal embryos, with an average length to width ratio of 1.4 ± 0.2 (**Fig. 3E**).

By contrast, only 1.1% of the embryos observed in GFP+ animals at day 4 post hatching appeared to be polynucleated (2/179 embryos imaged across 16 animals), with the remainder appearing to have normal morphology (**Fig. 3 C–E**). Both apparent polynucleated embryos in this genotype had two nuclei and may have been fertilized embryos in the middle of the first division. The average length to width ratio of all embryos from GFP+ animals is 1.7 ± 0.3 (P_adj_ = 3.25 e-18 vs GFP–).

By Day 6 post hatching, all the embryos in GFP– animals were polynucleated (169/169 embryos imaged across 11 animals), whereas 28% of GFP+ embryos were polynucleated (52/184 embryos imaged across 15 animals, **Fig. 3D**). As such, heterozygous animals also displayed phenotypes observed in the homozygous siblings but seemed to take longer to manifest. A larger fraction of the GFP+ embryos imaged at day six post hatching were also misshapen, with an average total length to width ratio of 1.5 ± 0.3, compared to 1.3 ± 0.2 for age matched homozygous controls (P_adj_ = 7.39e-11), and 1.7 ± 0.3 for GFP+ embryos imaged on day 4 post hatching (**Fig. 3E**). Interestingly, not all GFP+ animals displayed the polynucleated embryo phenotype. Five of the fifteen animals imaged (33%) contained no polynucleated embryos, indicating the phenotype is not fully penetrant in older heterozygotes.

### Mutants die by bursting at a young age

Estimates of the lifespan of wild-type (N2) adult hermaphrodite *C. elegans* in the laboratory range from 11.8 to 20 days [37]. The variability between published estimates is caused by spontaneous alleles that are rapidly fixed in lab stocks grown under non-selective growth conditions. To determine whether the microdeletion mutation reduces the life span of the worm, we again compared homozygous GFP– animals to genetically matched GFP+ heterozygous siblings. We synchronized animals as arrested L1 larvae, then isolated individuals of each genotype by following pharyngeal GFP fluorescence. Eight days after hatching, 64% of GFP+ animals and 58% of GFP-animals survived, suggesting that there is no large difference in lifespan between heterozygous animals and homozygous siblings. (**Fig. 4**). Strikingly, in both genotypes, a significant number of animals died by bursting (**Fig. 4, SMovie 1**), although the phenotype appears to be more prevalent in homozygous GFP– animals. By day eight, 35% of all GFP– animals died by bursting, while only 11% of GFP+ animals died by bursting. The cause of death for the remainder of the animals was not determined.

**Figure 4.**
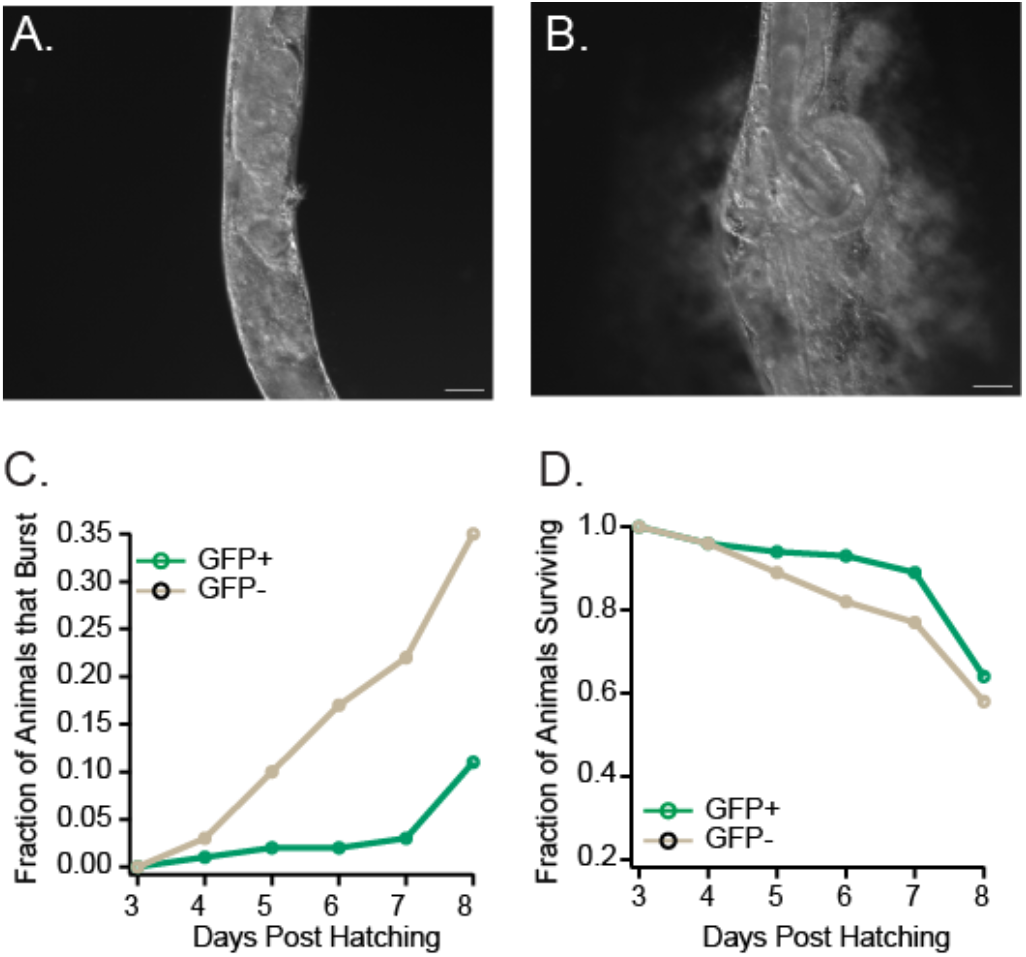
Microdeletion *sprDf1* mutants are prone to death by bursting. DIC micrographs of a GFP– *sprDf1* homozygous worm observed while bursting are shown. The full movie is available as SMovie 1. **A**. This frame corresponds to time zero, the beginning of the movie. **B**. This frame corresponds to the same animal, 4 minutes and 57 seconds later. The scale bars in both panels represent a distance of 20 microns. **C**. The fraction of animals that died by bursting as a function of days posthatching is shown with GFP+ heterozygotes in green and GFP– homozygous siblings in beige. The data shown represent the sum of at least three independent synchronizations, with N ranging from 26–166 animals per genotype per day. **D**. The fraction of animals surviving is represented as in panel C irrespective of cause of death.

### The microdeletion mutation causes a squashed vulva phenotype

We were intrigued by the bursting phenotype and wondered if vulva morphogenesis defects might contribute to the bursting phenotype. Notably, the microdeletion eliminates the *sqv-6* gene, which had previously been identified in a forward genetic screen for the squashed vulva phenotype. Previously characterized *sqv-6(n2845)* mutants display a striking decrease in the vulval lumen volume during the larval L4 stage and defects in formation of the UTSE valve between the uterus and the vulva [38]. To assess whether the microdeletion induces a similar phenotype, we imaged the vulva of L4 homozygous and heterozygous animals on day three post synchronization. Consistent with loss of *sqv-6*, GFP– homozygotes had a mean luminal width at half maximal height of μm 0.55 ± 0.8 μM, while the luminal width of GFP+ siblings was five times larger on average (2.74 ± 1 μm, P_ks_ = 9.5e-22, **Fig. 5A–C**). GFP+ animals displayed a broad range of vulval widths, with 9.2% of animals imaged having a squashed vulva equal to or narrower than the mean of GFP– animals. The full distributions are displayed in **figure 5C**.

**Figure 5.**
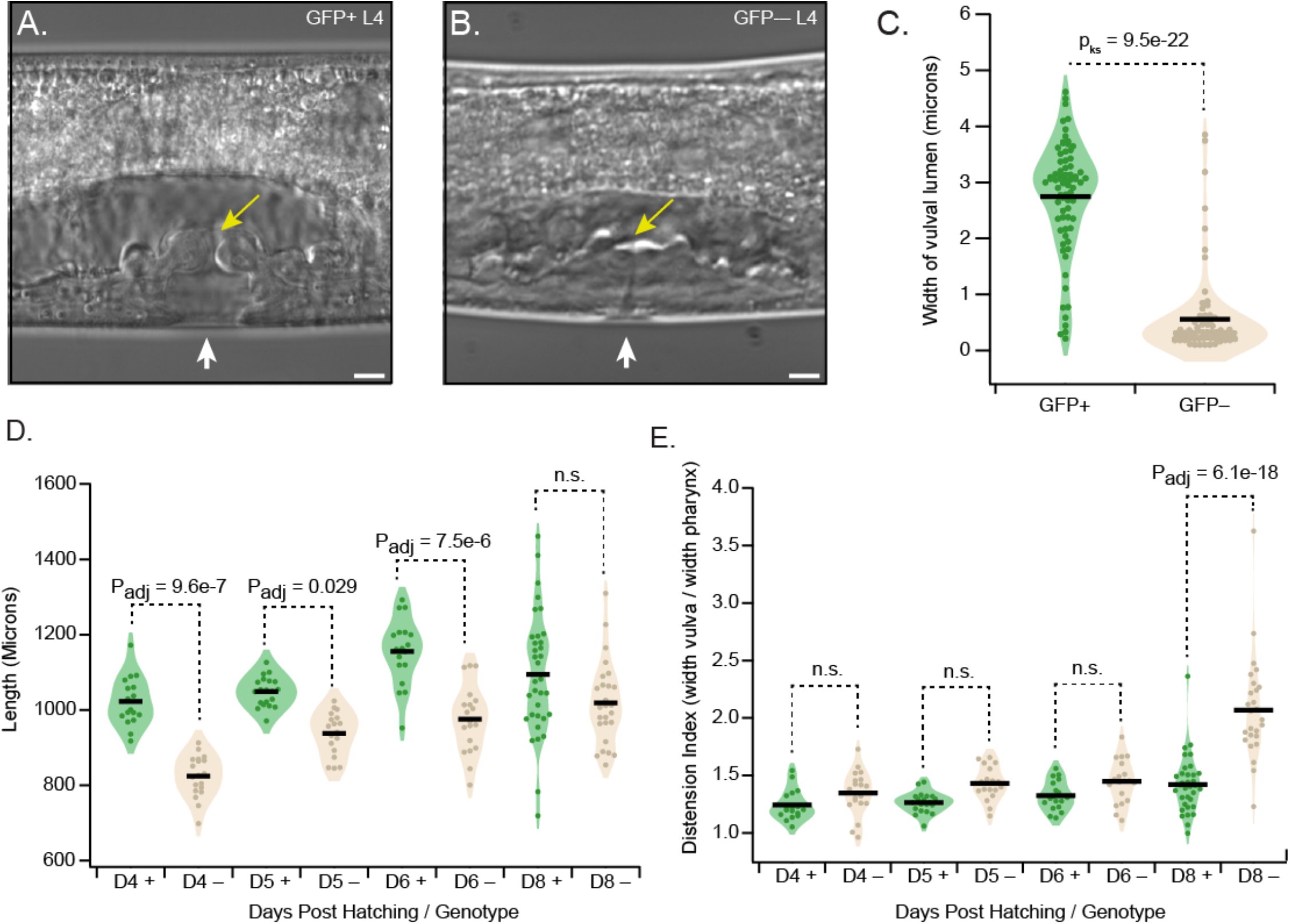
*sprDf1* homozygotes have a squashed vulva and are shorter and wider than their heterozygous siblings. **A**. DIC micrograph of an L4 GFP+ heterozygote showing the developing vulva. The vulval lumen is indicated with a white arrow, and the UTSE valve which separates the uterus from the vulva is indicated by a yellow arrow. **B**. Equivalently aged L4 homozygous *sprDf1* worms display a dramatic reduction in the width of the vulval lumen (a *sqv* or *squashed vulva* phenotype, white arrow), and the UTSE valve does not form properly (yellow arrow). In panel A and B, the scale bar represents 5 microns. **C**. The distribution of luminal widths of the vulva is shown for homozygous animals versus heterozygous siblings. The p-value was calculated using a non-parametric Kolmogorov-Smirnov test. Each point represents the luminal width from a single animal, and black bars represent the mean. **D**. The overall length of GFP+ heterozygotes and GFP-homozygotes is shown as a function of days post hatching. Each point represents the length of a single animal. The black lines represent the mean. P-values were calculated using a one-way ANOVA with the Bonferroni post-hoc correction for multiple hypothesis testing. **E**. A distension index, defined as the width at the vulva divided by the width at the pharynx, was calculated for each genotype from images collected as a function of days post hatching. The distribution, the mean, and statistical significance calculations are represented as in panel D.

### Homozygous microdeletion mutants are short and become distended with age

Animals homozygous for the microdeletion also appeared both shorter and wider than their heterozygous siblings. To quantify these phenotypes, animals were synchronized as L1 larvae, clonally isolated, then mounted and imaged as a function of time post synchronization using DIC microscopy. The overall length, the width at the vulva, and width of the lower bulb of the pharynx were measured for approximately 20 animals of each genotype from day three through day eight post hatching. Between day three and day six, homozygous GFP– animals were 10-20% shorter than heterozygous controls (**Fig. 5D**, Day 4: 19%, P_adj_ = 7.1e-7, Day 5: 11%, P_adj_ = 0.026, Day 6: 15%, P_adj_ = 5.8e-6). The difference was less apparent as the animals aged and was not significant by day eight (Day 8: 7%, P_adj_ = 0.12), suggesting a slow or delayed growth phenotype. To quantify the apparent thickening of homozygous animals compared to heterozygous siblings, we divided the vulval width measurement by the width of the lower bulb of the pharynx to define a distension index that could be compared across siblings of both genotypes (**Fig. 5E**). This index revealed that homozygous GFP– animals were notably thicker than heterozygous siblings by day eight post synchronization (Distension Index Day 8 = 2.1 GFP+, 1.4 GFP-, P_adj_ = 6.1e-18), but not at younger ages. This suggests the thickening manifests quickly in older animals and is likely caused by the unregulated growth of the embryonic tissue in the uterus. Animals that survived longer than 8 days could not easily be mounted and imaged due to bursting while handling.

### Homozygous microdeletion mutants are uncoordinated

Another striking phenotype observed in animals homozygous for the microdeletion is strongly uncoordinated movement. Adult animals become progressively paralyzed as they age. Larval animals and young adults appear to move relatively normally, but older animals have difficulty moving their tail, and do not travel far on a standard NGM-agar plate (**Fig. 6A, SMovie 2**). To quantitate these phenotypes, we again synchronized WRM31 animals at the L1 stage by bleaching and hatching, then monitored the animals as a function of days post-hatching by filming GFP+ heterozygotes or GFP– homozygotes for 25 seconds on an unseeded NGM agar plate. Prior to filming, each animal was touched with a worm pick to stimulate movement. Animals were filmed for 25 seconds at a frame rate of 4 images per second. The films were analyzed using the wrMTrck plugin for the FIJI implementation of ImageJ to determine both the length of the path traveled and the average worm velocity [39, 40]. Movies were collected for both genotypes at day 5 and day 6 post hatching. Comparing only worms that could be tracked across the entire assay window, GFP+ worms traveled farther on average than their homozygous GFP– siblings (**Figure 6B, C**, Mean Path Length = 2600 ± 1200 μm Day 5 GFP+, 1100 ± 800 μm Day 5 GFP–, n=22 and 32 respectively, P_adj_ = 3.4e-6; 2600 ± 1500 μm Day 6 GFP+, 920 ± 400 μm Day 6 GFP-, n=15 and 18 respectively, P_adj_ = 5.0e-5). Comparing all worms imaged, GFP+ worms traveled at a higher rate of speed than GFP-siblings at both time points (Mean Velocity = 120 ± 50 μm/sec Day 5 GFP+, 40 ± 30 μm/sec Day 5 GFP–, n=40 and 42, respectively, P_adj_ = 7.8e-10; 130 ± 70 μm/sec Day 6 GFP+, 50 ± 40 um/sec Day 6 GFP–, n=15 and 18, respectively, P_adj_ = 3.7e-8). Older animals could not easily be filmed due to their propensity to burst when handled.

**Figure 6.**
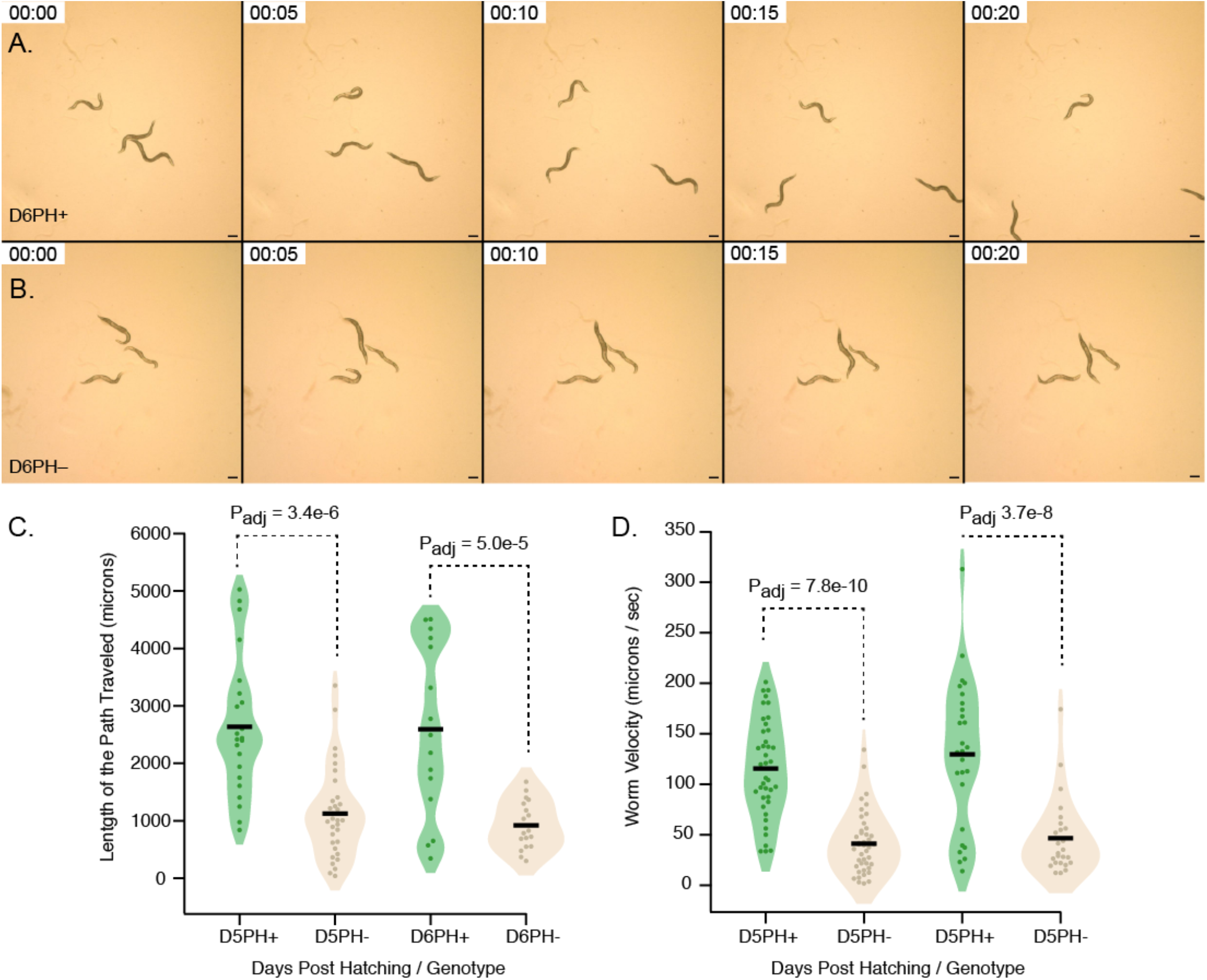
Homozygous microdeletion mutant animals are uncoordinated. **A**. and **B**. Still frames from movies collected for GFP+ heterozygous worms (top row) and GFP– homozygous worms (bottom row). The full movie can be seen in SMovie 2. The time stamps indicate time post initiation of filming, and the scale bar represents 200 microns. **C**. The total length of the path traveled is shown for each worm that could be tracked for 25 seconds. Each point represents individual animals (green = GFP+, beige = GFP–) and black bars represent the mean. The P-values were calculated with a oneway ANOVA with Bonferroni post-hoc correction for multiple hypothesis testing. **D**. The average velocity (microns/second) of all worms was tracked across each frame. The data plotted and analyzed as in panel C.

## DISCUSSION

Microdeletions are an important class of genetic lesion responsible for a wide variety of human developmental syndromes [1]. They are caused by chromosomal deletions that affect multiple genes but are too small to be detected by normal karyotyping (< 5MB). They tend to arise spontaneously and can induce a wide variety of diseases, including 22q11DS (DiGeorge Syndrome), Prader-Willi syndrome, Neurofibromatosis type 1 and 2, and many others. The penetrance of the phenotypes in microdeletion syndromes is variable and can be influenced by the extent of the deletion, dosage compensation, and genetic background effects. Moreover, as 90% of the cases of 22q11DS are spontaneous [1, 5], it remains unclear if there are certain genetic backgrounds that predispose individuals to generate the microdeletion.

How simultaneous haploinsufficiency of many adjacent genes leads to strong yet variable phenotypes is not well understood and has not been directly assessed in a simple model organism. We have recovered a 255KB multi-locus microdeletion (*spr1Df1)* in *C. elegans* that is similar to the microdeletion alleles in a variety of human developmental diseases. Mutants that harbor this microdeletion are viable and fertile as heterozygotes, survive to adulthood, yet are sterile as homozygotes, and display multiple striking phenotypes in both genotypes. Reminiscent of the microdeletion syndromes, heterozygotes display widely disparate phenotypic penetrance, even among siblings from the same brood. We suggest that this strain could provide a useful model to investigate the complex relationships between pathways that are simultaneously disrupted due to microdeletion-like alleles. And while *C. elegans* is evolutionarily distant from humans, and there is no clear synteny between this microdeletion and any specific microdeletion syndrome in humans, the robust genetics, rapid gene disruption technology, and simple RNAi screening enables characterization of the nonlinear relationships between genotype and phenotype at a level that is intractable in higher eukaryotes.

### Genes that potentially contribute death by bursting

The 255KB *sprDf1* microdeletion removes 32 adjacent protein coding genes, including 16 that have not been studied and 16 more where we have some information about their function from the literature. Of these, genetic screens have identified phenotype-inducing alleles in six of them, four of which also show phenotypes in RNAi screens. These genes are *rnst-2, nra-4, ergo-1, unc-34, sqv-6*, and *fmo-5*. No single gene deletion can account for all of the phenotypes observed in *spr1Df1* homozygotes, nor the variable penetrance. As such, it is likely that the phenotypes are caused by the simultaneous disruption of multiple genes.

Both heterozygous and homozygous animals are prone to premature death by bursting. We suggest bursting is likely caused by a combination of factors. First, vulval morphogenesis defects are observed in both genotypes, although they are more prevalent in GFP– homozygotes. These defects are likely caused by loss-of-function of the *sqv-6* gene, which was previously recovered in a screen for the squashed vulva (*sqv*) phenotype. The *sqv-6(n2845)* allele causes a vulval morphogenesis phenotype identical to that shown in **figure 5B**. This nonsense allele results in loss of 42 amino acids on the C-terminus of *C. elegans* sole homolog of XYLT1, a polypeptide xylosyltransferase required for proteoglycan synthesis. On its own, the *n2845* allele does not cause bursting. The presence of polynucleated embryos in the microdeletion mutant could also be attributable to disruption of the *sqv-6* gene. The *sqv-6(n2845)* mutation leads to embryonic arrest at the one-cell stage without nuclear division. However, polynucleated embryos have been observed in published RNAi studies of *sqv-6*, and are also observed in other *sqv* gene mutants involved in proteoglycan synthesis [41]. The unregulated growth of these embryos into a uterine tumor, and subsequent bursting of the animals, was not observed in either the *n2845* mutant or in *sqv-6* RNAi animals. As such, it is likely that that disruption of additional genes in the microdeletion further contributes to the observed phenotypes.

Another gene that potentially contributes is *nas-32*. This gene encodes an astacin-like metalloprotease endopeptidase involved in molting. Published studies describing loss-of-function mutations and/or RNAi phenotypes of related family members revealed defects in the molting cycle, suggesting a role for these enzymes in modifying the architecture of the cuticle [42, 43]. It could be that unregulated growth of embryonic tissue in the uterus, coupled to defects in vulval morphology leading to uterine retention, and defects in the cuticle combine to cause premature death by bursting. Other genes lost in the microdeletion could also contribute to the phenotype, including additional factors potentially involved in proteoglyan carbohydrate remodeling (*clec-203*, Y50D4B.4) and additional proteases related to cathepsin B (*cpr-5* and *cpr-8*). These hypotheses remain to be directly tested.

### Genes that potentially contribute to the reduced brood and shortened lifespan phenotypes

The *rnst-2* gene encodes a homolog of the ribosomal RNA T2 endonuclease required for autophagy of ribosomes in the lysosome. Loss of *rnst-2* activity leads to partially penetrant maternal effect embryonic lethality (20-30%), larval arrest (50%), and a shortened lifespan [44]. Animals deficient in *rnst-2* accumulate ribosomal RNA and have larger than normal lysosomes. While *rnst-2* mutants share some features of the *sprDf1* phenotype, including shortened lifespan, there is no evidence that *rnst-2* mutants die by bursting, and the embryonic lethality phenotype of the *sprDf1* homozygotes is much more penetrant (100%) than *rnst-2* null mutants alone. It is possible that loss of *rnst-2* is responsible for premature death not caused by bursting in the microdeletion mutants, but it remains possible that other genes lost in the microdeletion also contribute to lifespan.

The *nra-4* gene encodes a transmembrane domain protein that associates with nicotinic acetylcholine receptors in the endoplasmic reticulum and regulates their assembly in synapses [45]. It is homologous to nodal modulator (NOMO), an agonist of TGFβ signalling pathways in vertebrate embryos. Hypomorphic alleles of this gene cause a variety defects in *C. elegans*, including mild resistance to aldicarb, levamisole, and nicotine, morphology defects at synapses, as well as a reduced brood size. RNAi of *nra-4* revealed a very low penetrance embryonic lethality phenotype (0.83%) as well. The mild phenotypes reported for this gene suggest that it alone does not account for the strong phenotypes reported above for the *sprDf1* microdeletion, although it is possible that loss of *nra-4* contributes to the reduced brood observed.

The *ergo-1* gene encodes an Argonaute family protein that associates with a subpopulation of 26G endogenous siRNAs that appears to play a role in silencing pseudogenes, recently duplicated genes, and long noncoding RNAs [46]. Loss of *ergo-1* mutants enhance RNAi phenotypes, but otherwise display no other physiological phenotype, and as such loss of the *ergo-1* gene is unlikely to contribute to the striking phenotypes of *sprDf1* described above.

### Genes that potentially contribute to the uncoordinated phenotype

The strong uncoordinated movement / tail paralysis phenotype observed in *sprDf1* homozygotes is almost certainly due to loss of the *unc-34* gene. Loss of function mutations in *unc-34* display a severe and fully penetrant uncoordinated phenotype characterized by paralysis, withered tails, and/or dramatic reduction in mobility relative to wild type animals [47-49]. UNC-34 encodes the lone homolog of Enabled / VASP in the *C. elegans* genome. Ena / VASP proteins promote the assembly of F-actin filaments in a variety of contexts [50]. In *C. elegans*, loss of *unc-34* disrupts the migration of several neuronal cell types, including CAN, PVQ, HLN, AVM, VD and DD neurons [47, 49].

In addition to *unc-34*, the *sprDf1* microdeletion also removes several uncharacterized genes that are expressed or enriched in neurons, including K10C9.7, K10C9.1, Y50D4B.2, and Y50D4B.1. The microdeletion also disrupts *flp-34*, which encodes a neuropeptide involved in olfactory learning processes [51]. We cannot rule out that loss of these genes contributes to the uncoordinated phenotype. Nor can we rule out that the uncoordinated movement / paralysis could be secondary to the unregulated growth of uterine tumor tissue, which occupies much of the body cavity.

### How did the microdeletion allele arise?

There are at least two potential explanations for how the microdeletion allele arose during our experiment. It is possible that one or more double strand DNA breaks were induced by Cas9 in our attempt to target the *mex-5* allele on chromosome IV. To assess the likelihood of this possibility, we used the program Cas-OFFinder to map potential off-target cleavage sites within the region removed in the microdeletion [52]. This analysis identified three weak off target sites within the deleted region, each of which contains five mismatches to the guides with a maximal bulge of 2 nucleotides. None are near the precise breakpoints. We suggest that it is unlikely that any of these sites was efficiently targeted by the injected Cas9-guide RNA complex. But we cannot rule it out.

The second possibility is that the allele arose spontaneously during our experiment prior to balancing. Indeed, there is evidence that this region of chromosome V may be prone to rearrangements. An overlapping segment of the N2 Bristol strain chromosome V (V:1,105,377–1,274,227) was found to be translocated to chromosome II in the divergent Hawaiian strain (CB4856) [53]. In microdeletion diseases, large deletions typically arise during meiosis through non-allelic homologous recombination, which can give rise to both duplications and deletions through mispairing of sister chromatids. No obvious homology is observed around the breakpoints of the *sprDf1* allele, suggesting that the microdeletion allele may not have arisen by this classic mechanism. However, extended microsatellite repeat sequences are located within a few hundred base pairs of both breakpoint sites [54]. An extended octonucleotide repeat sequence ATGCCTAC is found in 27 copies upstream of the left breakpoint, and 26 copies of a hexanucleotide repeat sequence CTAAGC are found upstream of the right breakpoint. Microsatellite repeat sequences are prone to expansion, mutation, and double strand breaks caused by strand reannealing difficulties during DNA replication [55, 56]. We suspect double stranded DNA breaks induced by difficulty in replicating the microsatellite repeats account for the large deletion.

Our work reinforces the importance of validating the genotype-phenotype relationship for alleles derived during CRISPR mutagenesis experiments. Directed sequencing methods that focus exclusively on predicted off target mutations could miss spontaneous mutations at unrelated loci, leading to misinterpretation. Given the low cost, we suggest routine re-sequencing of the *C. elegans* genome for alleles derived in this species. Out-crossing candidate mutants, and generation of multiple lines where possible, can help mitigate this risk, and potentially reveal new mutants with interesting properties.

## MATERIALS AND METHODS

### Strains and Nematode Culture

All strains used in this study were maintained by growing the animals at room temperature on *E. coli* OP50 seeded NGM plates under standard conditions [57]. The nematode strains used in this study are listed in **Table 1**. The WRM31 was generated by injecting N2 worms with an RNP-mixture containing recombinant *Streptococcus pyogenes* Cas9, a guide RNA targeting *mex-5* (protospacer: UGGAAUCAAACCAUGUGAUA), a second guide RNA targeting *dpy-10* (protospacer: GCUACCAUAGGCACCACGAG), tracr RNA (IDT, Coralville, IA), and a PCR product derived from amplifying DNA encoding the CX10 mutant of mex-5 [33] synthesized as a gBlock (IDT, Coralville, IA) and topo-cloned into a vector using the Invitrogen kit (CX10 gBlock: AAA AAA AAG GAT CCG GAC CCT TTC CGC CGA ATT ATA AAA CCC GCC TCT GCA AAA ACT TTG CAC GTG GCG GCA CGG GCT TTT GCG ATA TGG GCG CAC GTT GTA AGT TCG CCC ACG GCC TGA AGG AAC TCC GTG CTA CGG ACG CAC CAG CCC GTT ACC CGA ATA ACA AAT ACA AAA CGA AAC TGT GTA AAA ATT TTG CGC GTG GTG GGA CAG GCT TTT GCC CAT ATG GCT TGC GTT GCG AGT TTG TAC ACC CGA CCG ATA AAG AAT AAT GAC TCG AGA AAA AAA A). Primer sequences are listed in **STable 3**.

**Table 1:**

| Strain ID | Genotype |
| --- | --- |
| N2 | Wild Type (Bristol natural isolate) |
| VC362 | *unc-5(e53) IV/nT1 [qls51] (IV;V); dpy-11(e224) V/nT1 [qls51] (IV;V)* |
| CB4856 | Wild Type (Hawaiian natural isolate) |
| WRM31 | *sprDf1 (V)/nT1 [qls51] (IV;V)* |

Candidate mutants were balanced by crossing fertile siblings of animals bearing phenotypic and sterile worms with VC362 (*unc-5(e53)* IV/nT1[qIs51] (IV;V); *dpy-11(e224)* V/nT1[qIs51] (IV;V)). F2 animals were isolated, and those that produced only GFP positive wild type-appearing progeny and GFP– uterine tumor-bearing progeny were propagated. To outcross the balanced WRM31 strain, N2 males were crossed with VC362 hermaphrodites, Pharyngeal GFP positive F1 male progeny were subsequently crossed with WRM31 GFP positive hermaphrodites. The F2 progeny that were GFP positive were singled and allowed to self-propagate, and the presence of the uterine tumor phenotype scored in pharyngeal GFP negative animals. The crosses were repeated 2 more times, for a total of a 3X outcrossing of the WRM31 strain. The genotype of WRM31 was confirmed by PCR and whole genome sequencing as described in **Figure 2**, using the methods outlined below. Note that the outcrossed WRM31 strain does not harbor a mutation in the *mex-5* or *dpy-10* loci.

### SNP Mapping

Recombination mapping experiments were performed by crossing heterozygous WRM31 hermaphrodites with CB4856 (Hawaiian) males. Hermaphrodite F1 progeny that were WT in appearance and lacked GFP were singled to individual plates and allowed to self-fertilize. F2 progeny that were confirmed to display the uterine tumor phenotype were singled and then lysed in lysis buffer (30 mM Tris pH=8, 8 mM EDTA, 100 mM NaCl, 0.7% NP-40, 0.7% Tween-20 + proteinase K) prior to use in single worm PCR reactions. Lysates were frozen at -80°C for at least 10 min, then incubated at 65°C for 1 hour and 95°C for 15 min prior to genotyping PCR using the primer sets described by Jorgensen and colleagues [36]. Each PCR product was digested with the DraI restriction enzyme, then run on a 2% agarose gel to visualize products indicative of recombination or lack thereof. The experiments were repeated across multiple animals to determine the frequency of recombination between the mutant and each marker on the Hawaiian strain.

### Whole Genome Sequencing

Genomic DNA from strains N2, VC362, and WRM31 were prepared using the Qiagen Gentra Puregene Core Kit A (Germantown, MD) according to the manufacturer’s instructions. One 60 mm plate of starved worms was used as source material. The quality and concentration of the genomic DNA prep was confirmed using a Qubit fluorometer (ThermoFisher, Waltham, MA). The recovered genomic DNA was sent to Novogene (UC Davis, CA) who prepared libraries, sequenced the genome to an average depth of greater than 15X, and returned mapped data to the reference *C. elegans* genome. Single nucleotide polymorphisms, insertions and deletions, copy number variations, and structural variants were identified in all three strains relative to the reference genome. We further refined this list to identify candidate mutations that were 1) covered by the nT1[qIs51] balancer, 2) located on chromosome V to the left of snp_Y69A9L, and 3) unique to WRM31. The list can be found in **STable 2**. The presence or absence of these mutations was confirmed by PCR comparing GFP+ to GFP-animals. Primer sets used to evaluate candidate mutations are listed in **STable 3**. The identity of the breakpoint in the microdeletion was confirmed by Sanger sequencing of the PCR products using primers that flank the breakpoint.

### RNA interference

Knockdown by RNAi was performed via soaking N2 animals in double-stranded RNA corresponding to the genomic cDNA sequence of the *srj-38* gene. Total RNA was extracted from N2 animals with Trizol and phenol-chloroform prior to cDNA synthesis using RT-PCR with the Superscript III One Step RT-PCR system with Platinum Taq DNA polymerase kit (ThermoFisher Scientific cat #: 12574026). The resultant PCR product was used as a template for in vitro transcription (IVT) to produce dsRNA with the Ambion MEGAscript T7 in vitro transcription kit (ThermoFisher Scientific cat #: AM1333) following the manufacturer’s protocol. The dsRNA was extracted with phenol-chloroform extraction then precipitated with isopropanol. The primer sequences used to prepare the cDNA used in the IVT reactions are in listed in **STable 3**. RNAi by soaking was performed as previously described [58]. Briefly, arrested L1 animals were treated with 4-8 μg of purified dsRNA in soaking buffer at 20°C for 24 hours, then plated onto NGM plates seeded with *E. coli* OP50 and placed in the incubator. Once the animals reached adulthood, the presence or absence of a uterine tumor was scored by inspection using a stereo dissecting microscope.

### Brood Size, Hatch Rate, and Bursting Assays

WRM31 animals were maintained by picking pharyngeal GFP+ heterozygotes under a fluorescence dissecting microscope to limit recovery of animals that lost the balancer chromosome. WRM31 animals were synchronized by dissolving the worms in 20% alkaline hypochlorite solution (3 mL concentrated Clorox bleach, 3.75 mL 1M filtered sodium hydroxide, 8.25 mL filtered MilliQ H2O). Embryos recovered from gravid adults were harvested by centrifugation, washed extensively with M9, then left in M9 to hatch overnight. L1 larvae were plated on 60 mm OP50-seeded NGM agar plates the following day. Forty synchronized WRM31 GFP+ and GFP– animals were singled out into OP50 seeded NGM agar plates at the L3/L4 stage. The animals were transferred to fresh plates two days after singling and transferred to a new plate daily to prevent overcrowding. Prior to transfer, the animals were recorded as alive, burst, or dead. The number of the progeny were recorded after transferring worms. The following day, the number of embryos that hatched were recorded. The brood size, hatch rate, fraction burst, and fraction surviving were counted for eight days. At least three biological replicates were used per strain/condition.

### In Utero Imaging of Embryo Morphology

WRM31 animals were synchronized as described above. On day four, a subset of animals from each category were mounted in 1 mM levamisole (Sigma-Aldrich, BP212) and imaged using DIC optics on a Zeiss AxioObserver 7 with a 40x objective. Multiple images were collected at different focal planes to capture the dimensions of the embryos. This process was repeated two days later with worms from the same synchronization. The presence of polynucleated embryos / cytokinesis failure was determined by looking at the number of nuclei per cell per embryo in the uterus. The fraction of polynucleated embryos present was calculated by dividing the total number of polynucleated embryos by the total number of embryos present. The lengths of the embryos were measured by using the line tool in FIJI to draw a line between the vertices and measuring the major axis. The width of the embryos was determined the same way for the minor axis. The length/width ratio was calculated by dividing the two measurements. Older worms were not imaged due to increased chance of bursting before or during the imaging process. Statistical significance in the differences observed between GFP+ and GFP-siblings was determined using a one-way ANOVA with Bonferroni post-hoc correction for multiple hypothesis testing.

### Squashed Vulva Imaging

WRM31 animals were synchronized as described above. Two days after plating the arrested L1 larvae on OP50 seeded NGM agar plates, animals at the L4 stage were isolated, assessed for the presence or absence of pharyngeal GFP, then mounted in 1 mM levamisole and imaged with DIC optics on a Zeiss AxioObserver microscope using a 20x objective. The width of the vulval lumen at the midpoint of its height was measured with the line tool in FIJI as described above. Statistical significance in the differences observed between GFP+ and GFP-siblings was determined using the nonparametric Komolgorov-Smirnov Test.

### Worm Size Measurements

Synchronized L1 WRM31 animals were plated on OP50-seeded NGM agar plates. Animals were imaged on day four, five, six, and eight post-hatching in 1 mM levamisole using a 5x DIC objective on a Zeiss Axio-Observer microscope. The presence or absence of pharyngeal GFP was determined at the same time by imaging GFP fluorescence. The length of each worm (in microns) was determined using the segmented line tool in ImageJ to draw a line from the tip to the tail through the center line of each animal. The width of each animal was measured using the line tool in FIJI both at the vulva and at the lower bulb of the pharynx. A distention index was calculated by dividing the width at the vulva by the width at the pharynx to normalize for overall size of the animal. Statistical significance in the differences observed between GFP+ and GFP-siblings was determined using a one-way ANOVA with Bonferroni post-hoc correction for multiple hypothesis testing.

### Worm Velocity and Path Length Assays

WRM 31 animals were synchronized via bleaching as described above. Two days after plating L1 larvae (three days post hatching), worms were separated based on presence of GFP in the pharynx. On day five and day six post hatching, worms were picked onto unseeded plates and filmed using a 5X objective on a Zeiss Discovery V20 stereo microscope fitted with a Diagnostic Instruments 18.2 Color Mosaic camera. The zoom was 15x. The worms were stimulated with a pick prior to the images being taken. Images were collected with SPOT software. The movies consisted of 100 frames, taken 0.25 seconds apart, resulting in a 25 second long four frame per second movie. The movies were analyzed using the FIJI plug-in wrMTrck [39, 40]. Prior to using wrMTrck, the backgrounds were removed using the FIJI tools Z-project and Image Calculator functions, and the image thresholds were calculated with the Otsu algorithm. The wrMTrck outputs (length, average speed) were converted from pixels to microns by using a stage micrometer to determine the number of pixels per micron at 15x (0.2 pixels per micron), and then using the conversion on the outputs.

## ACKNOWLEDGEMENTS

The authors thank members of the Ryder and Massi labs for helpful comments on this manuscript. We thank Reyyan Bulut and Hannah Snell for assistance with imaging during the early stages of this work, Dr. Nick Rhind and Dr. Oliver Rando for sharing equipment and for helpful advice, and Wen Chen and Dr. Paul Sternberg for providing a list of physically mapped multi-locus deletion alleles curated by Wormbase. Some strains were provided by the *Caenorhabditis* Genetics Center, which is funded by NIH Office of Research Infrastructure Programs (P40 OD010440). This work was supported by NIH Grants R01GM117008 and R01GM139316 to S.P.R. and F.M.

## AUTHOR CONTRIBUTIONS

KA, DT, FM, and SR designed the CRISPR mutagenesis experiments, KA and DT generated the reagents necessary to perform the CRISPR mutagenesis. KA, PC, SN, and SR designed and performed experiments to identify and evaluate the mutation. SR wrote the manuscript in consultation with all authors.

## SUPPLEMENTARY MOVIE LEGENDS

**SMovie 1**. Bursting Phenotype. The movie shows a time lapse of a GFP– *sprDf1* homozygous animal bursting. The time stamp is listed in the upper left hand corner, the video has been speed up to a rate of 4 frames per second for convenience.

**SMovie 2**. Uncoordinated Phenotype. The left panel shows three GFP+ *sprDf1* heterozygous animals crawling on an unseeded NGM agar plate after being touched with a worm pick. The right panel shows the same except with three GFP– *sprDf1* homozygotes. The time stamp is listed in the upper right hand corner. The frame rate is 4 frames per second and the movies are presented in real time.

## SUPPLEMENTARY FIGURES AND LEGENDS

**Supplemental Figure 1.**
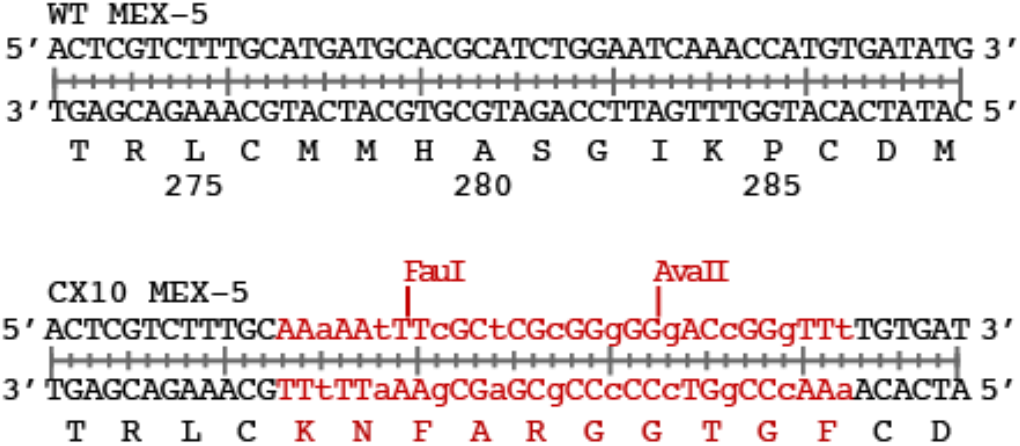
Sequence of the recoded region of MEX-5 in the CX10 mutant (in red). The position of the restriction enzymes used to detect the knock in variant is annotated. Lower case letters denote the wobble position of the recoded amino acids.

